# Daily ultrastructural remodeling of clock neurons

**DOI:** 10.1101/2024.11.06.622332

**Authors:** JI Ispizua, M Rodríguez-Caron, FJ Tassara, KY Kim, C Insussarry Perkins, M Barzi, C Carpio-Romero, MF Vasquez, CN Hansen, J Gargiulo, E Rosato, H de la Iglesia, MH Ellisman, MF Ceriani

## Abstract

In *Drosophila,* about 250 clock neurons in the brain form a network that orchestrates circadian rhythmicity. Among them, eight small Lateral ventral Neurons (s-LNvs) play a critical role, synchronizing the circadian ensemble *via* the neuropeptide Pigment-Dispersing Factor (PDF). Moreover, their neurites show daily variations in morphology, PDF levels, synaptic markers and connectivity. This process, called circadian structural plasticity, is ill-defined at the subcellular level. Here, we present 3D volumes of the s-LNv terminals generated by Serial Block-face Scanning Electron Microscopy (SBEM) at three key time points, two hours before lights-ON, two hours after lights-ON, and two hours after lights-OFF. We report a reduction in the number of neuronal varicosities at night, which reflects (and probably regulates) the cycling of the components we found therein. Indeed, in the morning we observed more presynaptic sites and increased accumulation and release of dense core vesicles. These rhythms were paralleled by periodic changes in mitochondrial structure that suggest daily modulation of their activity. We propose that circadian plasticity of the functionally relevant structures within presynaptic varicosities cyclically modulates the influence of the s-LNvs on the clock network.

## Introduction

The selective pressure imposed by the day-night cycle through evolutionary time has favoured the emergence of circadian clocks. These are complex, endogenous mechanisms that align biochemical, physiological and behavioral functions to the external environmental cycles, thereby increasing fitness^1,2^. In the fruit fly *Drosophila melanogaster*, the circadian clock comprises a network of almost 250 brain neurons^3^. Among them, the small Lateral ventral Neurons (s-LNvs, four per hemisphere) express and release Pigment-Dispersing Factor (PDF), a neuropeptide^4^ that is necessary to maintain rhythmic behavior under constant laboratory (aka free-running) conditions^5–8^. Additionally, these neurons express short Neuropeptide F (sNPF), regulating feeding and sleep, and the inhibitory neurotransmitter glycine, which contributes to the synchronization of the circadian network^9–11^. The s-LNv neurites project to the dorsal protocerebrum where their terminals exhibit rhythmic and stereotyped changes in the arborization pattern^12^, being more elaborated in the morning and less at night. Indirect markers of neuronal activity, such as anti-PDF immunoreactivity^13^ and the abundance of BRP^short^-RFP (which labels presynaptic sites^14^) are also rhythmic and peak in the morning^15^. Indeed, those markers and the complexity of the axonal arbour increase and decrease upon forced depolarization and silencing, respectively, confirming the functionality of structural remodeling^16–18^.

In this work, we explored how these manifestations of rhythmic neuronal activity interconnect at the subcellular level, using quantitative volumetric electron microscopy (EM). We analyzed three volumes from three adult female flies, each representing a key moment during the structural remodeling cycle: *Zeitgeber* Time 22 (ZT22, two hours before lights-ON), ZT2 (two hours after lights-ON), and ZT14 (two hours after lights-OFF). We labelled the s-LNvs by expressing an EM- detectable marker in the mitochondria (mito-matrix APEX2^19^) and generated 3D volumes of the terminals using Serial Block-face Scanning Electron Microscopy (SBEM) and a semi-automated convolutional neural network (CNN) algorithm to aid 3D reconstruction. In the three volumes, we identified and quantitated dense core vesicles (DCVs, carrying neuropeptides), presynaptic sites (PSs, anatomical equivalents of the T bars of the larval neuromuscular junction, a model synapse^20^) and mitochondria. Strikingly, we observed that these elements are concentrated in varicosities, enlargements of the plasma membrane that appear like beads on a string under light microscopy^21^. We show that the DCVs accumulate and fuse to the plasma membrane mainly in the morning, and that the PSs are more abundant in the morning as well. Moreover, such changes are associated with a marked shift in mitochondrial shape and volume between early morning and night. Finally, the number of varicosities also cycles, being higher in the morning. Our data indicates that the varicosities function as more than a release site of DCVs as previously assumed^22^. We suggest they represent a unit through which the s-LNvs modulate their influence onto the circadian network across the day. Such an organization, where varicosities represent discrete, regulated functional units of neuronal transmission, would better align the release of neurotransmitters and neuropeptides in space and time and optimize the relative arrangement of the components for better metabolic support. In summary, we report ultrastructural evidence of a rhythmic change in the number and relative composition of varicosities in the s-LNvs, which hints to a daily assembly/disassembly cycle. We suggest it may represent an efficient strategy to orchestrate daily fluctuations in connectivity, and ensure rhythmic but coherent transmission of information to the circadian network by the s-LNvs.

## Results

### Mitochondrial labeling allows rapid segmentation of *Drosophila* neurons in 3D EM volumes

To investigate circadian plasticity at the ultrastructural level, we generated 80x80x100 µm 3D EM volumes containing the s-LNv terminals by SBEM. To guide the identification of the neurons of interest we took advantage of APEX2, an engineered pea peroxidase^19^. The enzyme catalyses the deposition of osmophilic diaminobenzidine (DAB), which is visualized as electron dense in EM images. To avoid obscuring cytoplasmic details of synaptic terminals we generated a novel fly reporter line targeting APEX2 to the mitochondrial matrix. In addition, to facilitate correlated light and electron microscopy (CLEM) for multimodal multiscale analyses that enable the identification of the region of interest, APEX2 was fused to the fluorescent protein mKO2. The resulting construct, UAS-mito::mKO2::APEX2, (hereafter referred to as mito::APEX2) was expressed with the *Pdf-*GAL4 driver specifically in the lateral ventral neurons (LNvs, small, s- and large, l-). Based on previous studies, we chose to process the time points showing the most dramatic structural changes in the terminals of the s-LNvs: early morning (ZT2), early night (ZT14) and late night (ZT22). Briefly, ZT2 shows maximum spread of the neuronal terminals, ZT14 shows minimum spread with no change in total length, and ZT22 shows retracted terminals^15^. One volume per time point was produced (**Figure 1a**). Female brains were used for future comparison to existing connectomes^23,24^. DAB-labelled mitochondria (arrowheads, **Figure 1b**) guided the manual segmentation of the images and the generation of models (**Figure 1c**) with IMOD, a program used for 3D reconstruction from tomographic or EM serial sections^25^.

**Figure 1.**
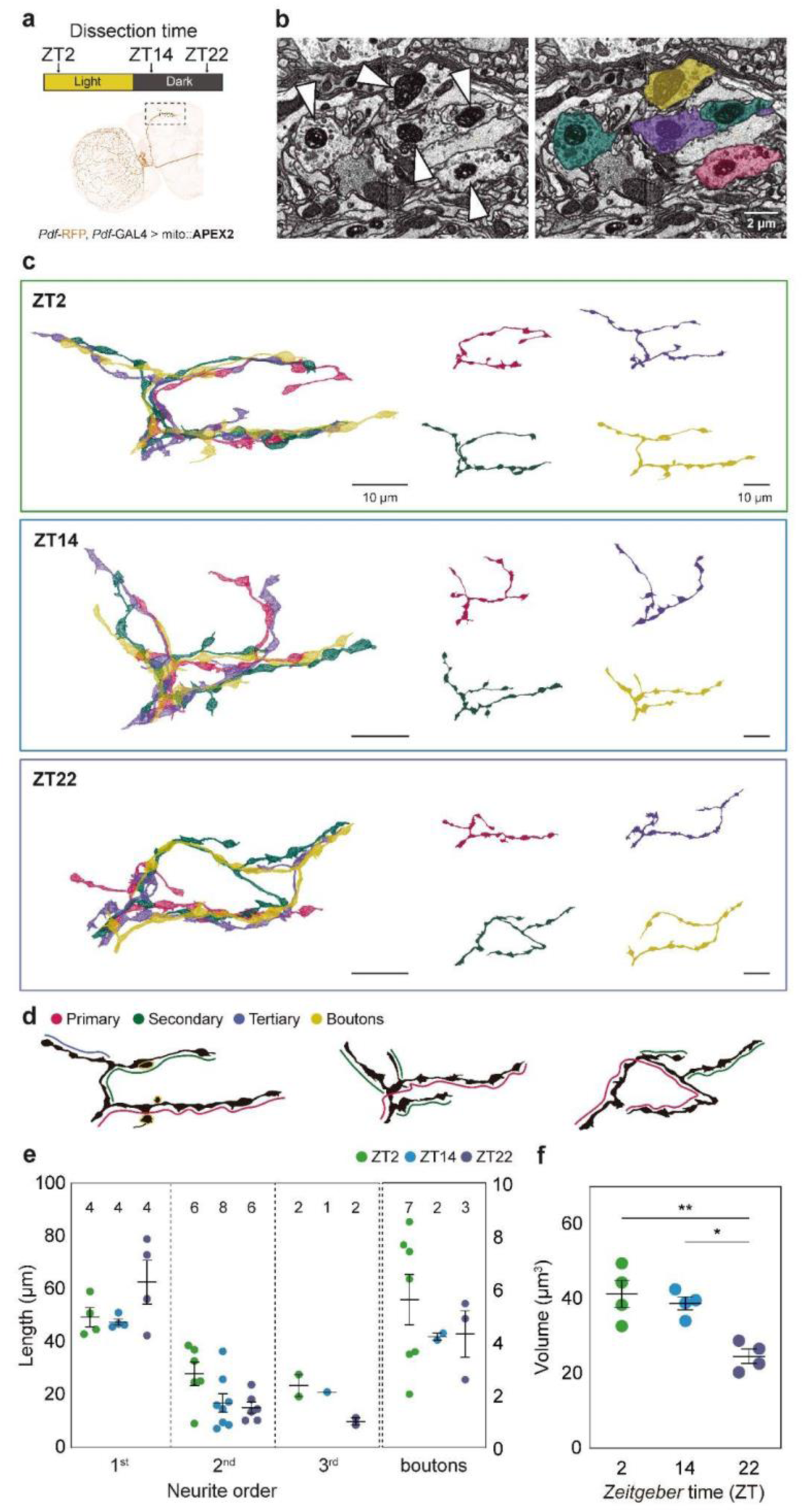
**Segmentation and 3D-models of the s-LNv terminals**. **a**, Schematic diagram of the experimental protocol. RFP expressed in PDF-positive neurons enabled the fluorescent identification of the s-LNv terminals for further processing. mito::APEX2 and DAB were employed to stain the mitochondria of the LNvs for SBEM. **b**, Labelled mitochondria (white arrowheads) were used to identify the s- LNvs terminals. **c**, 3D-models of the s-LNv terminals after manual segmentation showing them together (left) or individually (right) at each time point. **d**, Representative neurites from the ZT2, 14 and 22 volumes highlighting the segments defined as primary (magenta), secondary (green) or tertiary (purple) neurites and boutons (yellow). Primary neurites are defined as the longest projection extending from the fasciculated axon bundle, secondary neurites are those stemming from the primary ones, etc. Short protrusions including a terminal varicosity are labelled as boutons. No striking differences between individual neurites were observed at any given time point. **e**, The total number of neurites of each order is indicated at the top. The graph shows the quantitation of the neurite length according to their order (as defined in **d**). **f**, Volume of terminals/neuron per time point. In all graphs, error bars indicate the standard error of the mean (SEM). Asterisks indicate statistically significant differences: * p < 0.05, ** p < 0.01, *** p < 0.001. Non-significant differences are not shown. Details can be found in **Supplementary Table 3**.

A direct comparison of the segmented neurons with the corresponding confocal images obtained exploiting mKO expression provided compelling evidence that the former correspond to the s- LNvs (**Suppl. Figure 1**). All three volumes were manually segmented, resulting in the data presented here. In addition, using the data from ZT2 and ZT14, we trained a semantic classifier to recognize and segment cells containing labeled mitochondria and confirmed the reliability of the tool through the analysis of the volume collected at ZT22. Furthermore, an algorithm utilizing convolutional neural networks was developed for the segmentation of free and fused DCVs (**Suppl. Figure 2**). Collectively, they provide a pipeline that will make segmentation more readily accessible for the next generation of 3D-EM volumes.

The ability to identify individual neurons enabled a detailed assessment of specific neurites contributing to the complexity of the structure throughout the day. Primary neurites are processes that originate from the cell body, usually the longest, while secondary neurites originate from primary ones, etc (for a schematic diagram see **Figure 1d**). Interestingly, while the total number of primary, secondary and tertiary neurites remained relatively constant (**Figure 1e**) their length changed according to the time of day. Secondary and tertiary neurites tended to be longer at ZT2, which, together with the observation of an increased number of boutons (small protrusions with a terminal varicosity), would contribute to the more complex morphology of the early morning^15^. In addition, the total volume became significantly reduced late at night (**Figure 1f**). We confirmed that the varicosities distributed along the neurites function as *en passant* boutons, concentrating all the elements required for communication (mitochondria, DCVs, presynaptic sites), as already hinted in foundational fly TEM papers^26^. In the remainder of the manuscript, the results of our quantitative analysis of these volume EM samples are presented using varicosities as the focus of analysis.

#### Daily changes in vesicle fusion underlie differential neuropeptide release

DCVs carry neuropeptides from the soma to the neuronal terminals. PDF and sNPF are the main neuropeptides produced by the s-LNvs^27^, which among a variety of additional functions, synchronize activity in the clock neuron network and contribute to the regulation of sleep, respectively^4,11,28^. Using the manually segmented EM models, we identified both free DCVs and those that had entered the release path (fused DCVs, fDCVs). This analysis showed that there was a striking difference in the number of free and fused DCVs as a function of time of day (**Figure 2a-f**), which were found mainly within varicosities at any time (**Suppl. Figure 3d and g**). The total number of free DCVs was higher in the morning compared to ZT14 and 22 (**Suppl. Figure 3b-d**).

**Figure 2.**
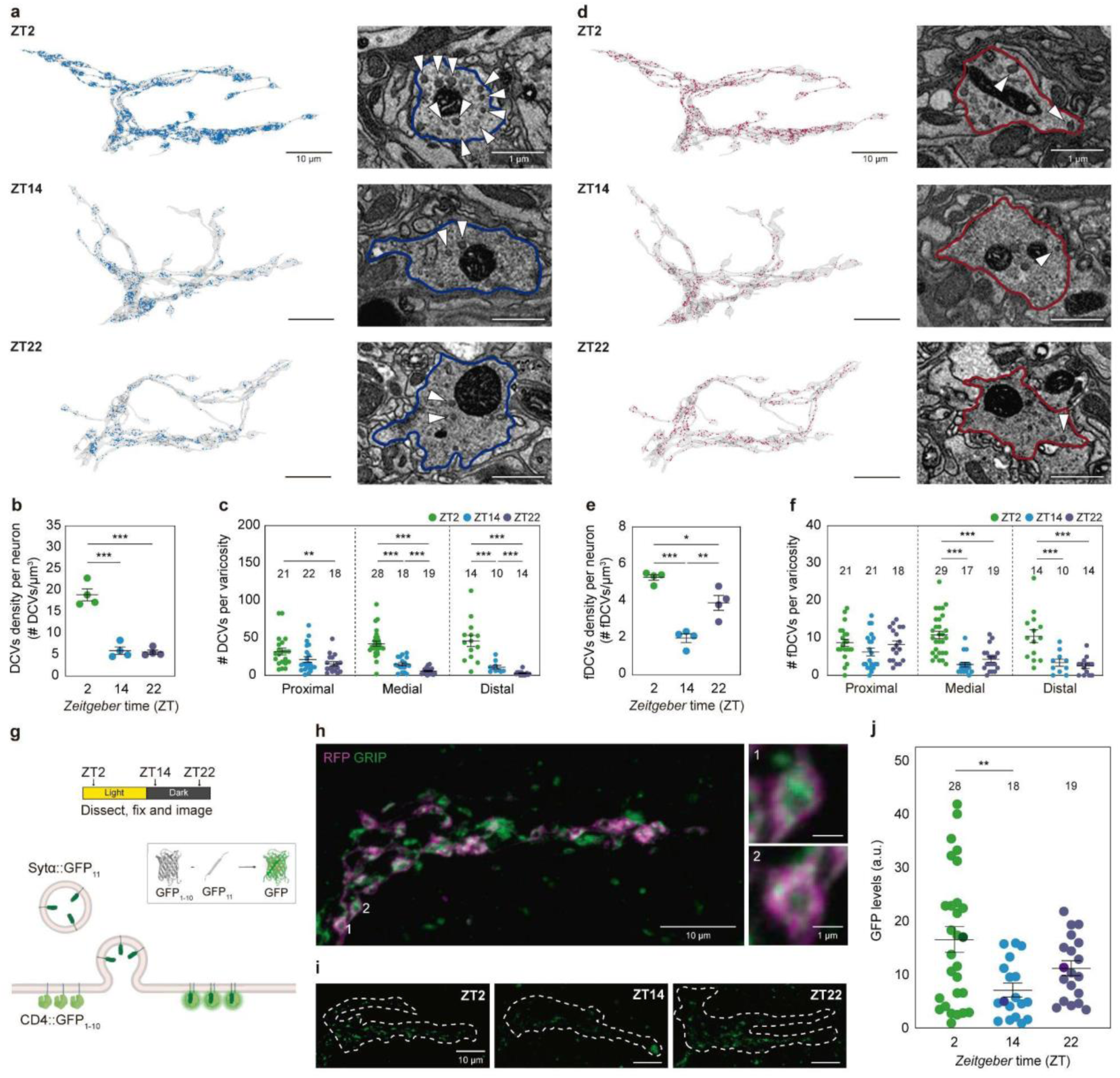
**DCVs availability and fusion change throughout the day**. **a, d**. 3D-models of neuronal terminals (on the left, in each panel) showing free dense core vesicles (DCVs, blue dots, **a**) and fusing dense core vesicles (fDCVs, red dots, **d**). Scale bars = 10 μm. Representative EM images (on the right, in each panel) show DCVs and fDCVs (white arrowheads). The plasma membrane is highlighted in blue and red in **a** and **d,** respectively. Scale bars = 1 μm. ZT2, ZT14, and ZT22 indicate the time points analyzed. **b**, **e**. Density per neuron (total number of vesicles/volume of the neuronal terminal) of free DCVs (**b**) and fusing fDCVs (**e**) at each time point. **c**, **f**, Total number of varicosities per condition examined is indicated above symbols. Graphs show the quantity (number of vesicles/varicosity) of free DCVs (**c**) and fusing fDCVs (**f**) per varicosity at each time point. Quantitation in proximal, medial and distal refers to the number of objects segmented in the initial, subsequent and terminal thirds of each neurite. Thirds are defined by the length of the longest neurite (See **Suppl.** Figure 3a for details). **g**, Schematic diagrams showing the experimental design (top) and the logic of the GFP reconstitution in the pre-synapse (GRIP) experiments (bottom). DCVs decorated with Sytα::GFP11 fused to the plasma membrane containing CD4::GFP1-10. GFP is reconstituted at release sites. **h**, s-LNv terminals expressing RFP (magenta) and GRIP (green). Maximum intensity projection of a Z stack of confocal images acquired with Airyscan modality. Scale bar = 10 μm. Individual varicosities are shown in insets 1 and 2; scale bars = 1 μm. **i**, Representative confocal images of GRIP (white hatched lines indicate the outline of the terminals). Figure 2**. DCVs availability and fusion change throughout the day**. **a, d**. 3D-models of neuronal terminals (on the left, in each panel) showing free dense core vesicles (DCVs, blue dots, **a**) and fusing dense core vesicles (fDCVs, red dots, **d**). Scale bars = 10 μm. Representative EM images (on the right, in each panel) show DCVs and fDCVs (white arrowheads). The plasma membrane is highlighted in blue and red in **a** and **d,** respectively. Scale bars = 1 μm. ZT2, ZT14, and ZT22 indicate the time points analyzed. **b**, **e**. Density per neuron (total number of vesicles/volume of the neuronal terminal) of free DCVs (**b**) and fusing fDCVs (**e**) at each time point. **c**, **f**, Total number of varicosities per condition examined is indicated above symbols. Graphs show the quantity (number of vesicles/varicosity) of free DCVs (**c**) and fusing fDCVs (**f**) per varicosity at each time point. Quantitation in proximal, medial and distal refers to the number of objects segmented in the initial, subsequent and terminal thirds of each neurite. Thirds are defined by the length of the longest neurite (See **Suppl.** Figure 3a for details). **g**, Schematic diagrams showing the experimental design (top) and the logic of the GFP reconstitution in the pre-synapse (GRIP) experiments (bottom). DCVs decorated with Sytα::GFP11 fused to the plasma membrane containing CD4::GFP1-10. GFP is reconstituted at release sites. **h**, s-LNv terminals expressing RFP (magenta) and GRIP (green). Maximum intensity projection of a Z stack of confocal images acquired with Airyscan modality. Scale bar = 10 μm. Individual varicosities are shown in insets 1 and 2; scale bars = 1 μm. **i**, Representative confocal images of GRIP (white hatched lines indicate the outline of the terminals).

Differences between the latter two times became significant in the medial and distal portion of the terminals (**Figure 2c**). The total number of fDCVs (**Suppl. Figure 3e-g**), albeit smaller than free DCVs, was also higher in the early morning. This suggests that DCV release is a function of the amount of DCVs available. Consistent with previous observations, fDCVs tend to be excluded from presynaptic densities^29^. The number of DCVs and fDCVs per terminal volume (DCV and fDCV density, **Figure 2b and e**) was higher at ZT2. However, at ZT22, fDCV density increased dramatically, probably because of the reduced volume of the terminals. Interestingly, this correlates with the growing excitability of the s-LNvs towards dawn^30^.

To further confirm daily regulation of neuropeptide release, we designed a new reporter of DCV fusion based on the recovery of fluorescence upon reconstitution of two split (non-fluorescent) GFP components. We called it GFP reconstitution in the pre-synapse (GRIP) because of its design. Briefly, one of the split GFP components was targeted to the lumen-facing side of the DCV membrane (*UAS-Synaptotagminα::GFP_11_*), while the other one was tethered to the extracellular portion of the transmembrane carrier protein CD4 (*UAS-CD4::GFP_1-10_* ^31^). The concomitant expression of both fragments in PDF neurons enabled the use of reconstituted GFP fluorescence as a proxy of DCV fusion (**Figure 2g**). As described for the EM dataset, most of the fluorescence was observed within the varicosities (**Figure 2h**). Notably, DCV fusion peaked early in the morning, reaching a minimum at the beginning of the night, and increased again towards the end of the night (**Figure 2i and j**), which is consistent with the observations made through SBEM in single volumes (**Figure 2e**). Altogether these results suggest time-of-day dependence of neuropeptide availability and release.

#### Presynaptic sites are more abundant in the morning

Synaptic sites are trans-cellular structures that organize fast neurotransmission between neurons. At the presynaptic site, many proteins organized in complex assemblies facilitate the docking of neurotransmitters-containing clear vesicles to the plasma membrane and their release after a depolarization event^32,33^. PSs were detected in the SBEM images (**Figure 3a**, model and arrowheads). They were reminiscent of T-bars and were surrounded by dense accumulations of unresolved structures that are likely clear vesicles. The majority of the synapses, if not all, were polyadic –contacting more than one postsynaptic neuron- and situated almost exclusively in varicosities and boutons (**Figure 3a**, right panel, asterisks, and **Suppl. Figure 3j**).

**Figure 3.**
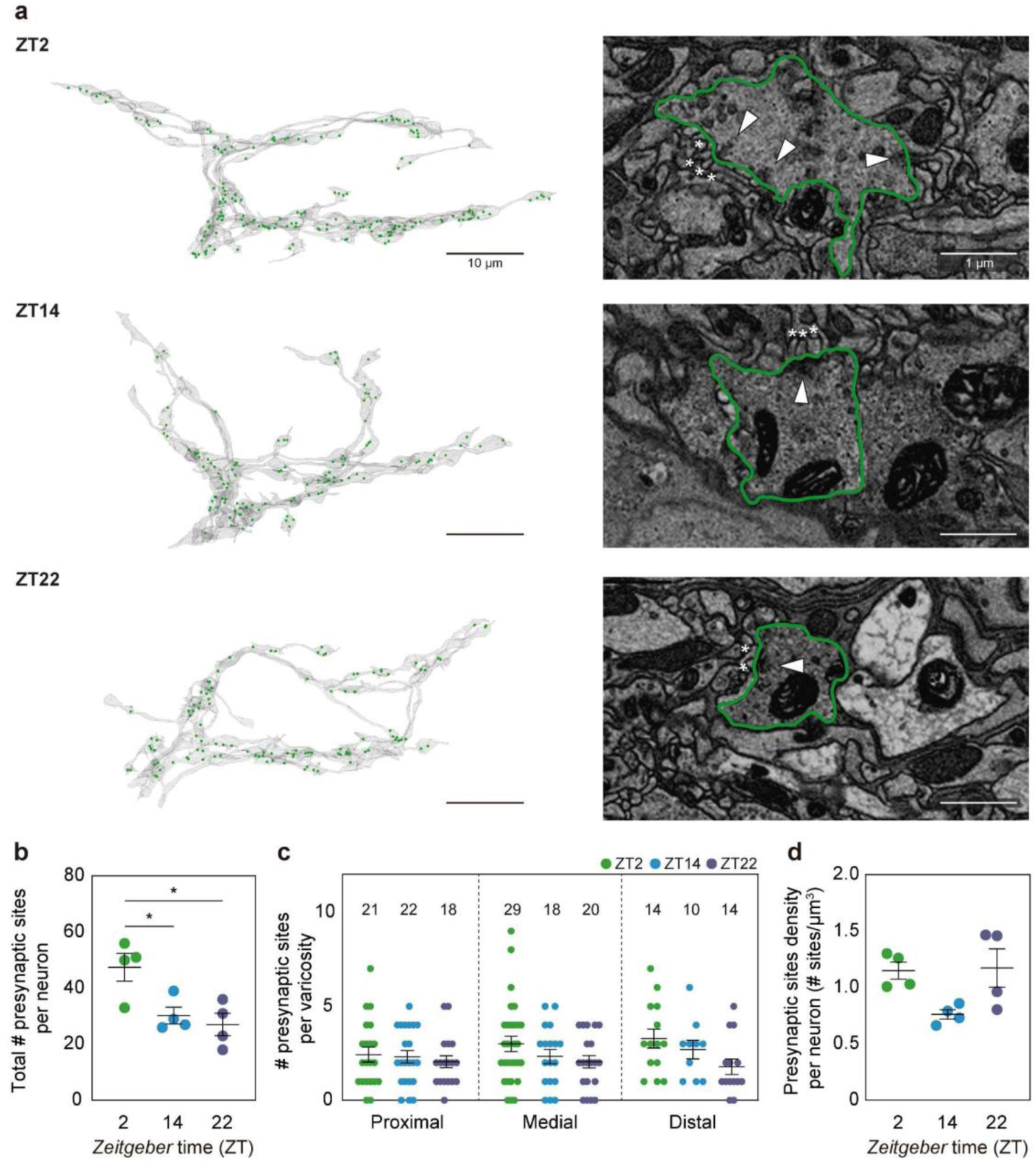
**Presynaptic sites are more abundant in the morning**. **a**, Segmented presynaptic sites/structures in 3D models of neuronal terminals (left, green dots, scale bars = 10 μm) and in representative EM images (right, PSs shown by white arrowheads, scale bars = 1 μm). Asterisks indicate putative post-synaptic structures. **b**, Total (including the objects located outside of varicosities) number of PSs at the different time points. **c**, Number of presynaptic sites per varicosity per region at each time point. The sample size (n, number of varicosities) is indicated above the symbols. **d**, Presynaptic sites density (total number of presynaptic sites/neuronal volume) at each time point. In all graphs, error bars indicate the standard error of the mean (SEM). Asterisks indicate statistically significant differences: * p < 0.05, ** p < 0.01, *** p < 0.001. Non-significant differences are not shown. Details can be found in **Supplementary Tables 1 and 3**.

Strikingly, the morning sample exhibited a two-fold increase in the total number of PSs in comparison to the volumes obtained at night (**Figure 3b**). Such difference likely results from a slightly higher number -yet non significant- of PSs per varicosity in the medial and distal segments (**Figure 3c, Suppl. Table 1**) together with a slightly higher number of varicosities at ZT2 (**Figure 5b**, right panel). When the number of presynaptic sites was normalized by the total volume of each neuron, the presynaptic density showed a decrease at the beginning of the night period, followed by an increase at ZT22, which is explained by the reduced volume of the terminals at that time (**Figure 3d**). Altogether these results confirm that differences in terminal complexity are associated with changes in presynaptic connectivity.

### Circadian structural plasticity correlates with changes in mitochondria shape and number

Mitochondria are dynamic organelles that exist in a fused-fragmented equilibrium. In extremely polarized cells, like neurons, they are usually found in close proximity to active zones^34^. They provide energy and serve as a Ca^2+^ reservoir^35^. In the s-LNv terminals, we identified marked time-of-day differences in the total number and morphology of mitochondria (**Figure 4a, Suppl. Figure 3k**), which were mostly detected within varicosities (**Suppl. Figure 3m**). No significant differences were observed in the total volume occupied by mitochondria (**Figure 4b**). Interestingly, at ZT2 we observed small and rounded mitochondria, usually more than one per varicosity (**Figure 4a**, right panel, arrowheads). However, at the end of the night we observed fused and elongated structures (**Figure 4a**), as indicated by a higher mitochondrial complexity index (MCI, **Figure 4c**). MCI approaches 1 as mitochondria become round (**Figure 4c**, top panel), whereas values >1 denote progressively complex and elongated shapes^36^. Our data suggests that during the four-hour interval between ZT22 and ZT2, the elongated mitochondria characteristic of late-night undergo fission.

**Figure 4.**
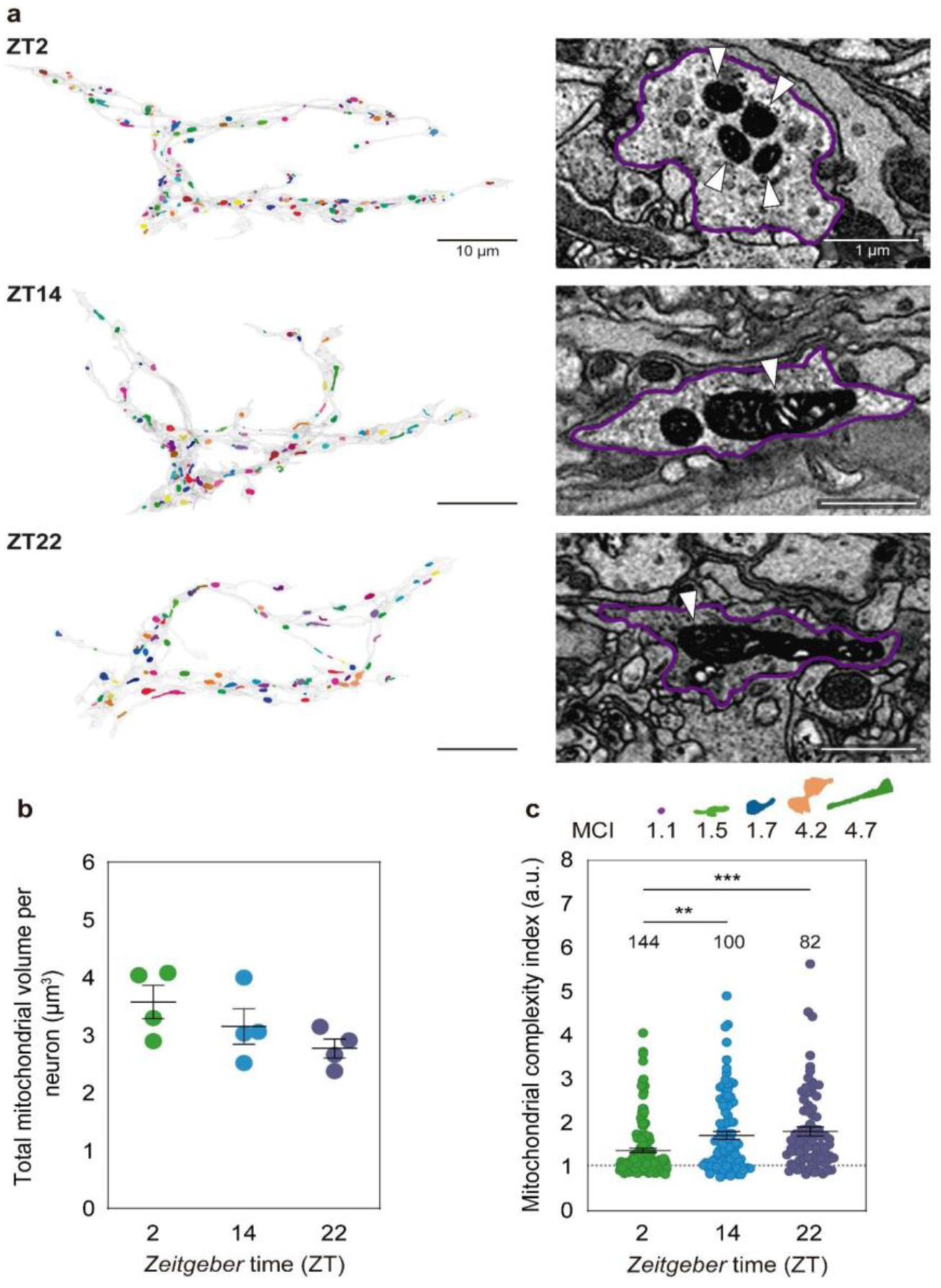
**Mitochondrial morphology changes along the day**. **a**, Segmented mitochondria (taken as individual objects, hence labelled in different colours) in modeled neurons (left, scale bars = 10 μm) and in representative EM images (right, scale bars = 1 μm) at each time point. Mitochondria are indicated with white arrowheads. **b**, Total mitochondrial volume per neuronal volume at the indicated time points. **c**, Mitochondrial complexity index (MCI) per time point. Top panel: an example of the shape of a mitochondrion along with the corresponding MCI. Bottom (graph): each symbol represents one mitochondrion. The total number of mitochondria (n) is indicated above the symbols. In all graphs, error bars indicate the standard error of the mean (SEM). Asterisks indicate statistically significant differences: * p < 0.05, ** p < 0.01, *** p < 0.001. Non-significant differences are not shown. Details can be found in **Supplementary Tables 2 and 3**.

**Figure 5.**
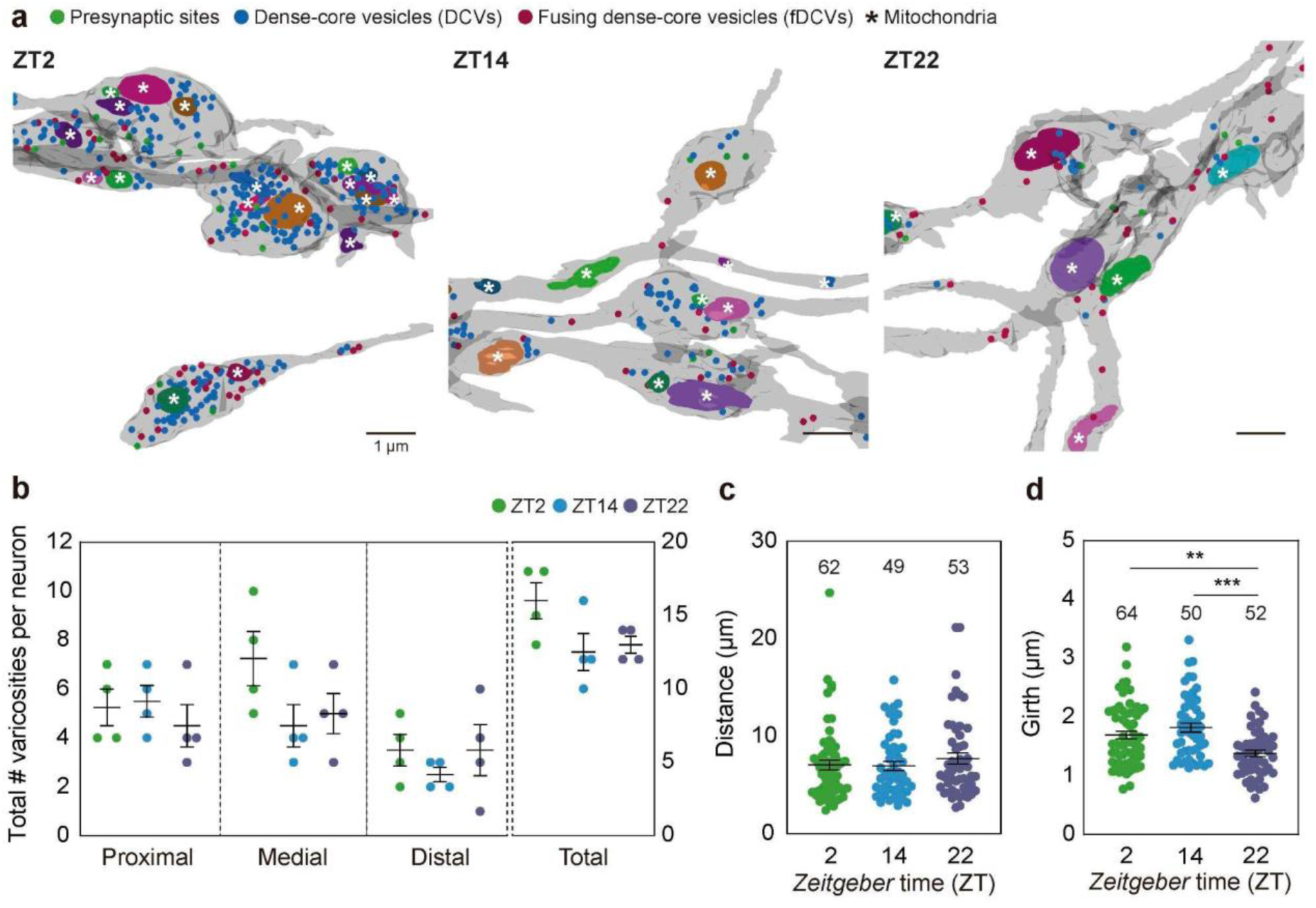
Varicosities show structural and functional plasticity. **a**, Representative model of the varicosities observed at different time points. Individual mitochondria are labelled in different colours. Scale bars = 1 μm. **b**, Total number of varicosities per neuron and region as explained in Fig. 2c. **c**, Distance between consecutive varicosities at each time point. The number of measurements (n) is indicated above the symbols. **d**, Varicosities girth at each time point. The number of analyzed varicosities (n) is indicated above the symbols. In all graphs, error bars indicate the standard error of the mean (SEM). Asterisks indicate statistically significant differences: * p < 0.05, ** p < 0.01, *** p < 0.001. Non-significant differences are not shown. Details can be found in **Supplementary Tables 2 and 3**.

#### Varicosities are plastic throughout the day

Thus far, we have shown that the varicosities of the s-LNv terminals are membrane enlargements with an approximate diameter of 1.8 µm at ZT2. They contain mitochondria and presynaptic sites and function as fusion hubs for neuropeptide-loaded DCVs (**Figure 5a** and **Suppl. Figure 3d, g, j, m**). In addition, we have observed that the total number of varicosities per neuron was higher in the morning than in the night samples. However, the differences were not evenly distributed, with the medial and distal parts of the volumes showing the largest variation (**Figure 5b**). Interestingly, while the distance distribution between varicosities remained stable at all time points (**Figure 5c**), suggesting that new varicosities are added at rather regular intervals, their size (girth) was reduced at ZT22 (**Figure 5d**).

Given that varicosities are a structural feature with a functional role, we propose that they represent the functional unit linking structural remodeling to functional plasticity.

## Discussion

### Generation of machine learning algorithms to guide the segmentation

Volumetric electron microscopy is an effective high resolution imaging method, generating extensive datasets from intricate three-dimensional samples. The application of technologies such as SBEM, serial section Transmission Electron Microscopy (ssTEM), and Focused Ion Beam Scanning Electron Microscopy (FIB-SEM) to neural tissue has led to the establishment of the field of *Drosophila* connectomics, pioneering the acquisition of functional insights from neuronal circuits^23,24^. In this study, we developed a combination of an established marker (mito::APEX2) with a new informatics interface to significantly reduce the processing load of 3D- EM data (**Suppl. Figure 2**). The mito::APEX2 marker, when used in concert with either the GAL4 or LexA expression systems, enables cell-specific targeting while circumventing the considerable opacity associated with membrane-bound APEX2, which has the potential to obscure synaptic sites and other fine ultrastructural features. Furthermore, it is possible to identify a cell without having to produce a volume that encompasses it entirely. This enables the reduction of the area of interest to specific zones of interest. Two manually segmented volumes (ZT2 and ZT14) were employed to train a semantic classification network to recognize labelled mitochondria. This resulted in the development of models for both the mitochondria and the neurons that contained them. Furthermore, an algorithm utilizing convolutional neural networks was developed for the additional segmentation of DCVs. Together, these advances provide a streamlined and accessible pipeline for targeted 3D-EM volume generation, expanding the potential of EM and connectomics to address specific biological questions.

### Circadian control of vesicle transport and release

Neurons use neuropeptides to convey messages that extend beyond individual synapses, producing broader cellular changes compared to the membrane potential shifts typically induced by classical neurotransmitters^37^. The s-LNvs produce at least two types of neuropeptides, both consisting of a short amino acid chain: PDF and sNPF. PDF reaches nearly 50% of the circadian neurons and some non-circadian, such as the lamina neurons^38^. It triggers an increase in cyclic AMP levels, which, in the circadian network, appears to modulate a wave of calcium oscillations sweeping across^28^. Conversely, sNPF seems to promote sleep consolidation by inhibiting the large ventrolateral neurons (l-LNvs), a group of clock neurons whose activation promotes wakefulness^39^. Both neuropeptides are transported within DCVs to the dorsal terminals of the s-LNvs, where they are captured until release^40^.

Since the early 2000s, it has been known that PDF levels cycle in a circadian manner, peaking in the morning within the dorsal terminals of the s-LNvs^13^. However, the dynamics of neuropeptide accumulation and release have only been analyzed indirectly, through fluorescent protein fusions with exogenous peptides^41^. Our work highlights the circadian nature of DCV accumulation within the dorsal terminals of the s-LNvs. We observe that free DCVs accumulate largely within the s-LNv varicosities, surrounding the mitochondria. Our data further shows that this accumulation decreases significantly at night, especially in the distal varicosities. A similar pattern is observed in DCVs undergoing fusion with the plasma membrane: fusion predominantly (although not always) occurs within the varicosities and exclusively outside the presynaptic densities, a pattern also described using fluorescent markers and expansion microscopy techniques^29^. Furthermore, we demonstrate that, similar to free DCVs, there are significantly more fusion events during the morning compared to nighttime, with a more pronounced effect in distal varicosities. The detection of fused DCVs at night indicates that, although most of the peptides are expected to be released from the s-LNv terminals at dawn —associated to the higher frequency of bursts of action potentials^30,42^— these neurons still release neuropeptides at night, despite the circadian variation in firing frequency^42–44^. This was further confirmed in acutely fixed brains through GRIP. The reconstituted (non-amplified) GFP signal was mainly restricted to varicosities, and the degree of fusion at the time points examined, judged by GFP intensity, closely resembles the EM data. In fact, a similar daily pattern was previously described employing a fluorescent DILP chimera at the s-LNv dorsal terminals, further confirming a temporal control in DCVs release^45^.

Interestingly, distal varicosities exhibited the greatest degree of plasticity in terms of the number of DCVs they could accommodate. Both the number of free and fused vesicles varied with a similar phase. This indicates that circadian modulation is likely to affect vesicle transport towards the dorsal projections, and that the number of vesicles fusing to the membrane may be contingent upon their availability, at least in part. A clear example supporting this hypothesis involves a s- LNv neuron observed at ZT22, where the primary neurite, after extending towards the midbrain, loops back into the axonal tract. This creates two non-contiguous varicosities, one proximal and one distal, separated (laterally) in space by less than 2 μm. If the release site were the primary determinant of the amount of vesicle fusion, both varicosities would be subjected to similar cues and behave similarly. However, the distal varicosity contains fewer DCVs and, consequently, significantly fewer vesicles fusing with the membrane compared to its proximal counterpart.

### The morning structure is more closely integrated with the network

Chemical synapses are functional structures that organize communication between neurons. The presynaptic neuron fuses vesicles filled with classical neurotransmitters (small molecules such as acetylcholine, glycine or glutamate), and the postsynaptic cell receives and interprets these neurotransmitters via membrane receptors, which induce downstream changes in membrane potential. In electron microscopy images of the *Drosophila* nervous system, presynaptic sites appear as clusters of clear vesicles organized around a T-bar structure formed by scaffold proteins, whereas postsynaptic sites only exhibit a slightly darker contrast in the membrane adjacent to the synaptic cleft^26,46^. Upon segmenting these structures in our volumes, we first observed that most synapses are polyadic, meaning that a single presynaptic density is associated with multiple postsynaptic neurons, consistent with previous observations^23,26^. Another interesting observation is that these synapses are mainly localized within varicosities, with the number of presynaptic sites per varicosity remaining largely stable throughout the day. There is a tendency, however, for varicosities to exhibit fewer presynaptic densities at night, especially in the distal regions. In any case, the morning organization displays approximately 50% more presynaptic sites than the two volumes taken at night time (**Figure 3b**), a finding that is consistent with previous reports based on fly brain samples examined using fluorescent protein labeling and 3D laser-scanning light microscopy^15^.

### Changes in mitochondrial structure imply variation in the signaling valence of the terminals and/or in energy demand

In most varicosities, vesicles and presynaptic sites are organized around one or more mitochondria. Given that both vesicle fusion and material transport rely on ATP, an organization into functional units appears to facilitate a localized ATP supply, thereby supporting essential communication processes. A recent study on mitochondrial morphology across the hemibrain demonstrated a correlation between mitochondrial shape and the identity of neuronal processes. Specifically, dendrites were observed to contain elongated mitochondria, while axons exhibited smaller, rounder ones^47^. This observation, coupled with prior evidence suggesting an increase in somatodendritic markers at the dorsal protocerebrum at night^46^, and even enhanced responsiveness to partner activation^49^, opens the possibility that the increase in mitochondrial complexity at night could be indicative of a transition from “axonal” to “dendritic” terminals.

An alternative, but not mutually exclusive, hypothesis is that changes in mitochondria morphology indicate a stress response. Just before dawn, the s-LNvs begin to fire at higher frequencies^30^, and at the same time they increase their presynaptic sites and show augmented recruitment and fusion of DCVs. Such an increased demand for energy could result in mitochondrial stress, resulting in the fragmented, rounded forms characteristic of ZT2. Conversely, at night, as neuronal activity decreases, mitochondria tend to fuse into larger, elongated structures with reduced total surface area. This configuration may enable collective evaluation of mitochondrial health and membrane potential equalization^50^, preparing the mitochondria for another round of metabolic stress at the start of the new day.

In mammals, some evidence suggests that the circadian clock controls the abundance and morphology of mitochondria by regulating biogenesis, fission/fusion and mitophagy^51^. Along those lines, the reduction in overall mitochondrial volume at the night onset could be indicative of clearance of damaged mitochondria through mitophagy. While electron microscopy images are unable to capture movement, observations of elongated mitochondria outside varicosities at night suggest that dismantling these functional units may contribute to the reduced complexity observed during this period.

### Varicosity changes link structure to function

Circadian structural plasticity was initially described through Z-axis projections derived from confocal microscopy images of brain samples collected at different time points. Thus, changes in complexity were characterized as variations in the number of crossing processes at increasing distance from the centroid using Sholl analysis^52^ and later on, defined by an image analysis algorithm^48^. Hypotheses regarding the types of subcellular changes underlying oscillations in complexity included the growth/disappearance of new neurites, the extension/retraction of existing neurites, and changes in the degree of fasciculation (i.e., proximity between neurites)^12^. However, distinguishing between these alternatives requires identifying individual neurites, an impossible task given the resolution of immunofluorescence coupled to confocal microscopy. While labeling individual neurons randomly could potentially circumvent this limitation, these stochastic labeling strategies not only exhibit low efficiency within the s-LNvs, but also do not consistently label the same neuron^53^.

Our data so far suggests that the number of neurites is similar across the three time points taken for analysis: in general, each neuron has one or two secondary neurites and only a few show tertiary neurites. Hence, changes in structural complexity would depend primarily on the growth/retraction (length) of individual neurites together with the generation of new varicosities and terminal boutons. Given the subtle nature of these changes, they are likely obscured by inherent structural differences across time points and the limited number of observations achieved with this level of resolution.

Interestingly, the high-resolution technique employed herein enabled the identification of additional changes. One such change was related to volume, whereby terminals occupied a greater space in the morning compared to the night, reminiscent of what was described in the L2 lamina neurons^54^. Another change was observed in the number of varicosities. This novel finding reveals a new dimension of circadian remodeling as it demonstrates that structural changes involve not only alterations in size or complexity, but also the addition of new, larger varicosities containing new synapses and peptide release sites. While most varicosities exhibited mitochondria, free and fused DCVs and PSs, a small fraction (about 5-6%) did not include any mitochondrion, while still containing free and fused DCVs and PSs. No varicosities contained exclusively PSs and, interestingly, no “empty” varicosities were found at any given time point.

This work provides evidence for daily changes in the size and volume of the s-LNv terminals, in addition to changes in the total number of varicosities, PSs, free and fused DCVs and mitochondria. These dramatic structural changes are likely to reflect functional differences, a claim further supported by the clear shift in the number and complexity of mitochondrial morphology, i.e. bioenergetics, across the day. The data collected thus far does not provide unequivocal evidence regarding the mechanisms that underlie the assembly and disassembly of varicosities. In the mammalian brain, high-frequency stimulation has been shown to induce axonal remodeling, whereby a transient enlargement of boutons is followed by a sustained widening of the axons, which in turn leads to functional changes^55^. Clock neurons in flies and mice exhibit circadian changes in membrane excitability, which in turn result in differential firing throughout the day^56–58^. While this phenomenon could contribute to the structural plasticity described herein^15^, it is tempting to speculate that there is a circadian component to membrane and/or vesicle biogenesis in addition to circadian transport (of presynaptic components, DCVs, etc) to support the extent of structural remodeling uncovered. This hypothesis has important implications for circadian rhythm disorders and disease.

## Materials and methods

### Fly rearing

Flies were grown and maintained at 25°C in vials containing standard cornmeal yeast agar medium under 12:12 h light:dark (LD) cycles. Adult-specific GAL4 expression for the GFP reconstitution experiments was accomplished through GeneSwitch^16^. GeneSwitch expression was induced transferring 2- to 5-day-old flies to vials containing food supplemented with RU486 (mifepristone, Sigma, USA) in 80% ethanol to a final concentration of 200 µg/ml for exactly 72 h prior to dissection. The *pdf*-GeneSwitch (*pdf*-GS) and the mito::mKO2::V5::APEX2 described in this manuscript) lines were generated in our laboratory. *Pdf*-GAL4 was obtained from the Bloomington Stock Center and *Pdf*-RFP was generously provided by J. Blau (NYU, USA). Stocks for GRIP were generated in the Rosato Lab and UAS- CD4::GFP_1- 10_ was provided by K. Scott (UC Berkeley, USA).

### Generation of transgenic lines

The mito::mKO2::V5::APEX2 sequence was provided by the Ellisman Lab^19^. A Kozak sequence was added upstream of the mito::MKO2::V5::APEX2 fragment. This sequence was synthesized and cloned into pJFRC81-10XUAS-IVS-Syn21-GFP-p10 (Addgene, USA) by GeneScript (GeneScript, USA), replacing the GFP coding region. Transgenic lines were produced by BestGene (BestGene, USA) by Phi31 site specific transformation into attP2 (third chromosome) or VKO2 (second chromosome) sites.

For GRIP analysis, the GFP_11_ fragment was fused through a flexible linker to the N-terminus end (pointing to the lumen of vesicles) of Synaptotagminα (Sytα). The DNA sequences encoding the fusion proteins were cloned into pUAST^59^ in the Xho I-Xba I sites. Transgenic lines were produced by the University of Cambridge Fly Facility (UK) using p-element transformation. Lines expressing split GFP components in the plasma membrane (UAS-CD4:GFP_1-10_) were already available and described^60^.

### GFP reconstitution in the pre-synapse (GRIP)

*Pdf*-GS > *Pdf*-RFP;GRIP flies were induced in RU for 72 h as described above. At ZT2, ZT14 and ZT22, males and females were briefly anesthetized on ice and their brains dissected in cold PBS under red light. Dissected brains were fixed in PFA 4% for 1h at room temperature in darkness. Samples were washed in cold PBS for 15 minutes (min) and then rinsed for 5 min. Finally, brains were mounted in Vectashield (Vector Laboratories, USA) with their dorsal surface upwards to allow access to the dorsal protocerebrum. Z-stacks of the s-LNv dorsal projections were taken immediately using a confocal microscope Zeiss LSM 880 (Zeiss, Germany) with a 40x water objective (N.A 1.2) employing the Zen software (Zeiss). The same objective and an Airyscan detector were employed to acquire the image in **Figure 2h**.

### Brain preparation for Correlative Light and Electron Microscopy

Briefly, 2- to 4-day old female flies expressing *Pdf-*GAL4>mito-APEX2;*Pdf-*RFP were ice-anesthetized and decapitated at the corresponding time points. Heads were fixed in 2.5% glutaraldehyde and 2% paraformaldehyde in 0.15 M cacodylate buffer (CB, pH 7.4) on ice for 1h. After removing the fixative, brains were dissected in fresh 0.15 M CB and washed 3 times for 10 min in the same buffer solution. To facilitate the penetration of the staining, an optic lobe was removed using fine forceps, leaving the adjacent region of interest undamaged. Brains were transferred to a coverslip and imaged at 20x, 40x, and 60x in an Olympus FluoView 1000 microscope (Olympus). Immediately after, brains were treated for 15 min with 20 mM glycine in 0.15 M CB on ice, to quench any unreacted glutaraldehyde. For the DAB reaction, a preincubation step was done in 2.5 mM DAB solution (25.24 mM stock in 0.1 M HCl) in 0.15 M CB for 15 min on ice. For staining, 0.03% H_2_O_2_ containing DAB solution was added to the brains on ice for 40 min. The staining buffer was washed off with ×5 washes using 0.15 M CB. Then, brains were individually placed inside scintillation vials into 1% OsO_4_ in 0.15 CB for 40 min at 4°C. Next, the OsO_4_ solution was removed and, without washing, a reduced solution of 1.5% potassium ferrocyanide in 0.15 M CB containing 2 mM CaCl_2_ was added. Brains were incubated for 1.5 h at 0°C and 30 minute RT. After thoroughly washing 3 x 10 min with ddH_2_O, brains were transferred to 0.5% aq. thiocarbohydrazide for 15 min at RT. The brains were washed 3 x 10 min on ddH_2_0 and stained with 2% aq. OsO_4_ for 30 min at RT. Brains were washed with ddH_2_O at RT 3 x 10 min and then stained with 0.5% aq. uranyl acetate for 30 min at RT. For the last staining step, after 3 x 10 min washes with ddH20 at RT, brains were incubated with 0.05% lead aspartate solution for 30 min at 60 °C and 1 h at RT. The brains were washed 3 x 10 min on ddH_2_0 and dehydrated in crescent percentages of ethanol and finally to a staining solution at specific temperatures and precise intervals, by using a freeze substitution chamber (Leica EM AFS2, Leica). Brains were individually transferred to small baskets and incubated as follows:

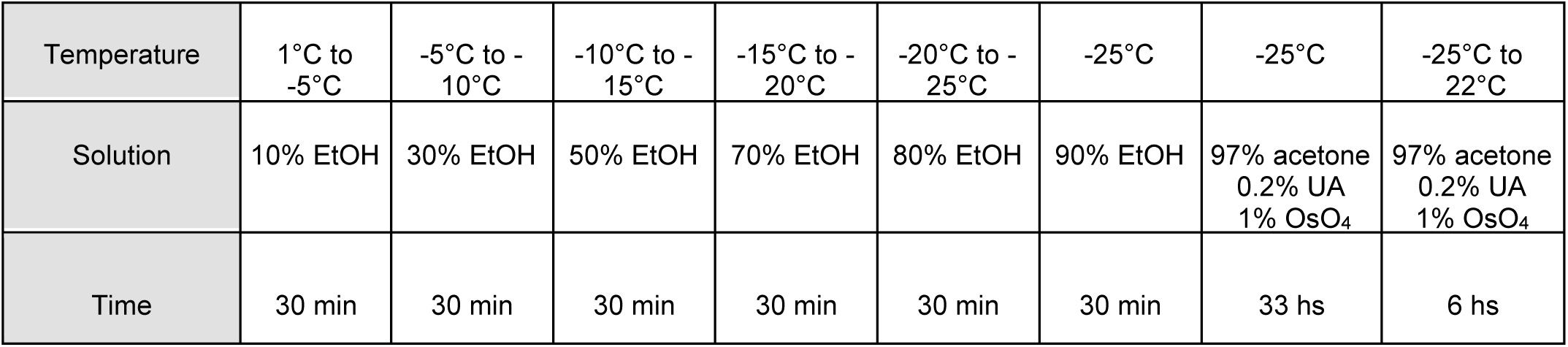

| Temperature | 1°C to -5°C | -5°C to -10°C | -10°C to -15°C | -15°C to -20°C | -20°C to -25°C | -25°C | -25°C | -25°C to 22°C |
| --- | --- | --- | --- | --- | --- | --- | --- | --- |
| Solution | 10% EtOH | 30% EtOH | 50% EtOH | 70% EtOH | 80% EtOH | 90% EtOH | 97% acetone<br>0.2% UA<br>1% OsO<sub>4</sub> | 97% acetone<br>0.2% UA<br>1% OsO<sub>4</sub> |
| Time | 30 min | 30 min | 30 min | 30 min | 30 min | 30 min | 33 hs | 6 hs |

Brains were then washed twice with dry acetone and placed into 50:50 Durcupan ACM:acetone overnight. Brains were transferred to 100% Durcupan resin overnight. Brains were then embedded in tin containers and left in an oven at 60°C for 72 hr. To proceed with Micro-computed tomography and SBEM, brains were carefully trimmed approximately ∼1 mm square pieces, leaving the minimum resin, and then mounted on Gatan SBEM specimen pins with conductive silver epoxy.

### Micro-Computed Tomography, definition of the region of interest and Serial Blockface Scanning Electron Microscopy

The Micro-Computed Tomography (Micro-CT) tilt series were collected using a Zeiss Xradia 510 Versa (Zeiss X-Ray Microscopy) operated at 80 kV (87 µA current) with a 20X magnification and 0.3836 µm voxel size for ZT2, 0.4687 µm voxel size for ZT14, and 0.4688 µm voxel size for ZT22. Micro-CT volumes were generated from a tilt series of projections using XMReconstructor (Xradia). Since no visible signal from the DAB deposition was seen in the Micro-CT, neuroanatomical features such as the alpha lobe of the mushroom bodies and the protocerebral bridge were used to correlate the computed tomography data to the confocal images. A 80*80*100 µm region of interest (ROI) containing the s-LNv terminals was defined for each timepoint. SBEM was accomplished using GeminiSEM 300 (Zeiss, Oberkochen, Germany) equipped with a Gatan 3View system and a focal nitrogen gas injection setup. This system allowed the application of nitrogen gas precisely over the blockface of the ROI during imaging with a high vacuum to maximize the SEM image resolution, as described in Deerink *et al* 2017^61^. Images were acquired in 2.5 kV accelerating voltage and 1 µs dwell time; Z step size was 50 nm; raster size was 16k × 14k for ZT2, 20k × 20k for ZT14 and 20k × 24k for ZT22 Z dimension was ∼900 to 1500 image samples. Volumes were collected using 90% nitrogen gas injection to samples under high vacuum. Once volumes were collected, the histograms for the slices throughout the volume stack were normalized to correct for drift in image intensity during acquisition. Digital micrograph files (.dm4) were normalized and then converted to MRC format. The stacks were converted to eight bits and volumes were manually traced for reconstruction using IMOD^25^. The s-LNvs were identified as neurons containing labeled mitochondria. They were segmented starting 2 microns below the first ramification of the first branching neuron of the bundle. DCVs were identified as dense, round structures of 90-200 nm. When connected to the plasma membrane, DCVs were marked as fusing DCVs. Synaptic sites were identified as a high electron density spot in the membrane with at least 2 post-synaptic partners, where a cloud of clear vesicles or a t-structure was deposited. These three elements were segmented as scattered objects. Mitochondria were recognized by shape, matrix pattern and high contrast due to DAB staining, and segmented as closed objects. Following the segmentation process, a single round of independent manual proofreading was conducted. The data pertaining to the total number of scattered objects, neuronal and mitochondrial volume, and surface area were extracted using the -*imodinfo* command. Quantitation was initially performed on a per-varicosity basis, with the varicosities themselves defined first, and then counting the objects within the model. For the purpose of quantitation, the primary neurite on each neuron was defined as the longest path from the point of origin of the structure. Subsequently, the relative position of each varicosity within a given neuron was normalized in relation to the length of the primary neurite of that neuron. Secondary neurites were defined as the longest protrusion emerging directly from a primary neurite. Tertiary neurites were defined as protrusions from secondary neurites. Boutons were defined as small sprouts containing one varicosity. The MCI was calculated using the equation described in ^36^:

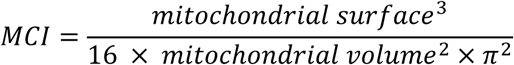

### Semi-automated segmentation

Three main neural networks support the tool’s segmentation tasks. First, a semantic segmentation model identifies s-LNv terminals by detecting stained mitochondria, which is then followed by a high-precision model that refines the boundaries of the mitochondria. A convolutional neural network (CNN) is then employed for the detection of vesicles within the boundaries of neurons. The segmentation models utilize Residual Attention U-Net on small patches (6.4 x 6.4 µm) to optimize efficiency, subsequently reassembling the patches to facilitate full volume segmentation. Neuron and mitochondrial annotations from previous IMOD data, refined in Python and ImageJ, provided ground truth for training on 4,150 images.

For vesicle detection, a "mobile classifier" (developed by PyTorch) uses a CNN^62^ with four layers, max-pooling, and fully connected layers on 250 x 250 nm patches to estimate the probability of vesicle presence. A second "refinement classifier" is employed to reduce the number of false positives, utilizing larger 500 x 500 nm patches to provide greater context around vesicle centers. The classifiers were trained and validated on a dataset comprising thousands of vesicle and non-vesicle images.

Finally, using py-clesperanto^63^ for GPU-accelerated 3D segmentation, each structure is labeled to facilitate metric calculations, like structure count and distribution within the volume.

### Quantitation and statistical analysis

Throughout the manuscript all the objects analyzed are the result of manual segmentation. The statistical analyses were performed using the R software, version 4.1.0.

*For the parameters displayed per neuron per time point (volume, DCVs, fDCVs and PSs density, total number and total number per region, mitochondrial volume and number of varicosities).* Datasets were analyzed using one-way ANOVA followed by t-test with Bonferroni corrections from the *emmeans* package (v. 1.10.5^64^).

*DCVs, fDCVs, and PSs per varicosity per region.* A model was applied to each neuronal region and object. Generalized linear mixed models with a Poisson distribution were fitted using the *lme4* package (v. 1.1-35.5^65^), with neurites considered as a random effect. In cases where the assumptions were not met (over-dispersion), a Conway–Maxwell–Poisson distribution was used employing the *glmmTMB* package (v. 1.1.10^66^). Then, an analysis of deviance (based on type II

Wald chi-square tests) was performed and Bonferroni corrections were applied to post hoc comparisons.

*GFP reconstitution in the pre-synapse.* GFP signal in the dorsal projections of s-LNvs was quantitated using ImageJ (NIH). Maximum intensity Z projection of *Pdf*-RFP signal was used to generate a mask of the region of interest (ROI). The ROI was used over a maximum intensity Z projection of GFP signal to quantitate mean intensity and then moved to the background to measure it. Background was subtracted to the terminal’s signal to get the final values. For statistical analysis, the fluorescence intensity in the terminals for each time point was modelled using a general linear mixed model, using the *nlme* package (v. 3.1-164^67^), in which the replica number was treated as a random variable. Furthermore, the model incorporates a parameter that allows the adjustment of the variance. One-way ANOVA was performed and subsequent analysis was conducted using the Bonferroni test.

*Mitochondrial complexity index (MCI)*. The mitochondrial complexity index was examined as a function of time point. A linear mixed model was fitted, and the variable was transformed using the natural logarithm. One-way ANOVA and a post hoc analysis with the Bonferroni test was conducted to compare between time points.

*Varicosity girth*. The data was fitted to a linear mixed model in which neurons were considered as a random variable. One-way ANOVA and *a posteriori* contrasts were performed with the Bonferroni test.

In all graphs, error bars indicate the standard error of the mean (SEM). Asterisks indicate statistically significant differences: * p < 0.05, ** p < 0.01, *** p < 0.001. Non-significant differences are not shown. Exact values for analyses and p values for contrasts are included in **Supplementary Tables 1-3**. The sample size (*n*) is indicated above symbols in graphs that have been statistically analyzed, except for the analyses per neuron, in which n=4 in all cases. The number of experiments performed is referred to as *N*. Outliers were defined with a first box plot exploration and confirmed with standardized residual analysis after modeling.

## Acknowledgements

We are grateful to the members of the Ceriani lab for their insightful discussions and to A. Liceri for providing fly food and assisting with fly work. We extend our thanks to M.I. Farías for her support in characterizing the APEX2 lines and to A.H. Rossi and A. Ross from the Microscopy and Bioimaging Facility at the Leloir Institute Foundation for their assistance and expertise in acquiring confocal microscopy images. We are indebted to I. Spiousas for his invaluable advice on statistical analyses. Our thanks also go to J. Blau for sharing the *Pdf*-RFP construct. Finally, we acknowledge the members of NCMIR for their training and guidance on EM sample preparation and analysis, as well as the Su Lab at UCSD for providing access to their fly facility.

## Consent for Publication

Not applicable.

## Availability of data and materials

All data generated or analyzed during this study are included in this published article and its supplementary information files. Any other data can be requested from the corresponding author.

## Author contributions

Experimental design: JII, FJT, MRC, MFC, ME, ER, HOI Data acquisition and analysis: JII, FJT, MRC, CCR, KK Statistical analyses: CCR

Generation of transgenic lines: JII, CNH, ER, MFC Algorithm development: FJT, CIP, MB

Manuscript writing and revision: JII, MRC, MFC, ER Review & editing of the final version: All authors

## Ethics declarations and consent to participate

Not applicable.

