## Supplementary figures and tables for "Daily ultrastructural remodeling of clock neurons"

Supplementary information Ispizua, Rodriguez-Caron et al.

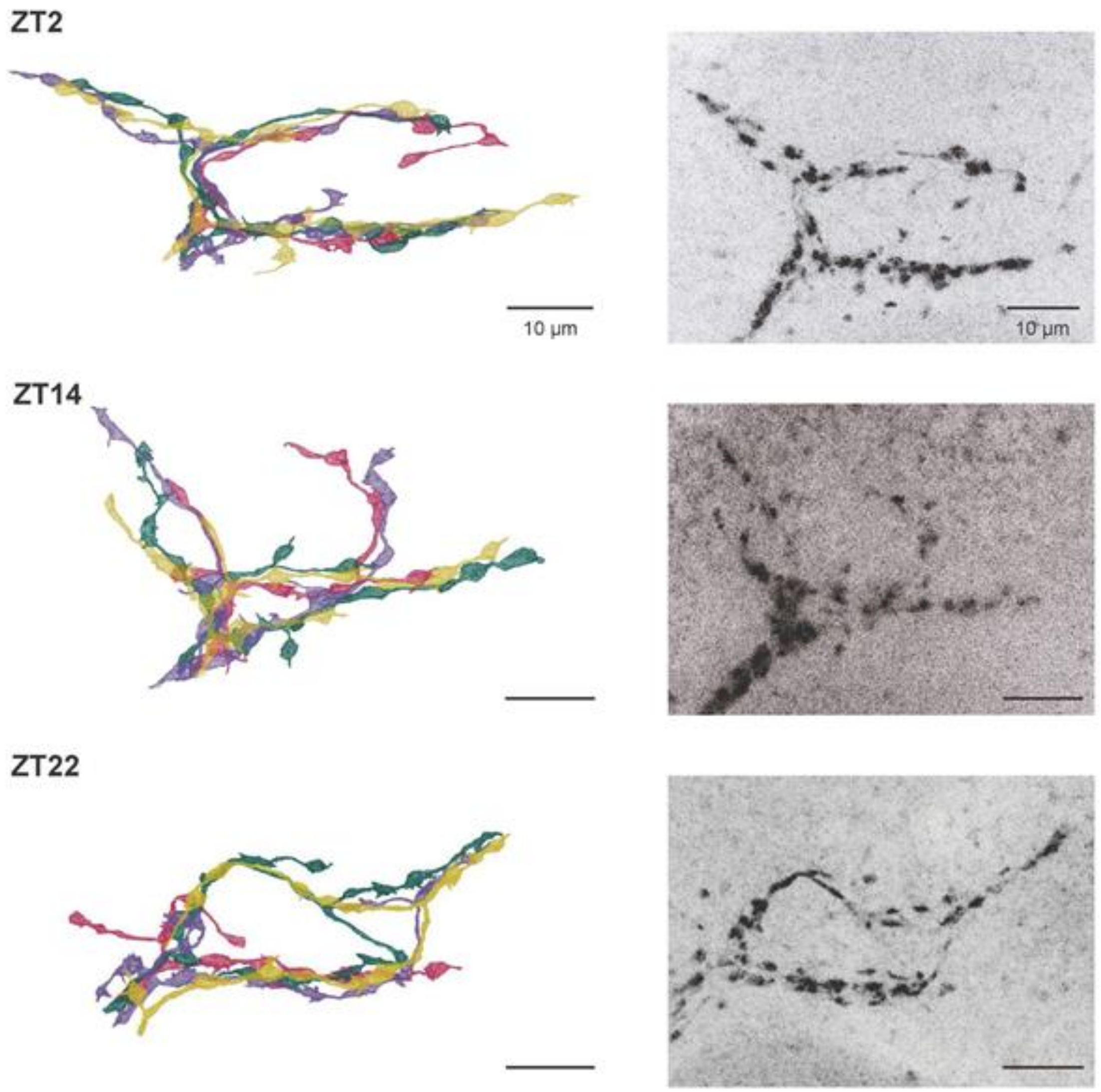


**Supplementary Figure 1. Comparison of the segmented models and the corresponding confocal images**. In the left column, the neuronal models for ZT2, 14 and 22 are shown. In the right column, Z-projections of confocal images obtained from the corresponding brains can be observed. Images were taken with a 63x 1.1 NA objective.

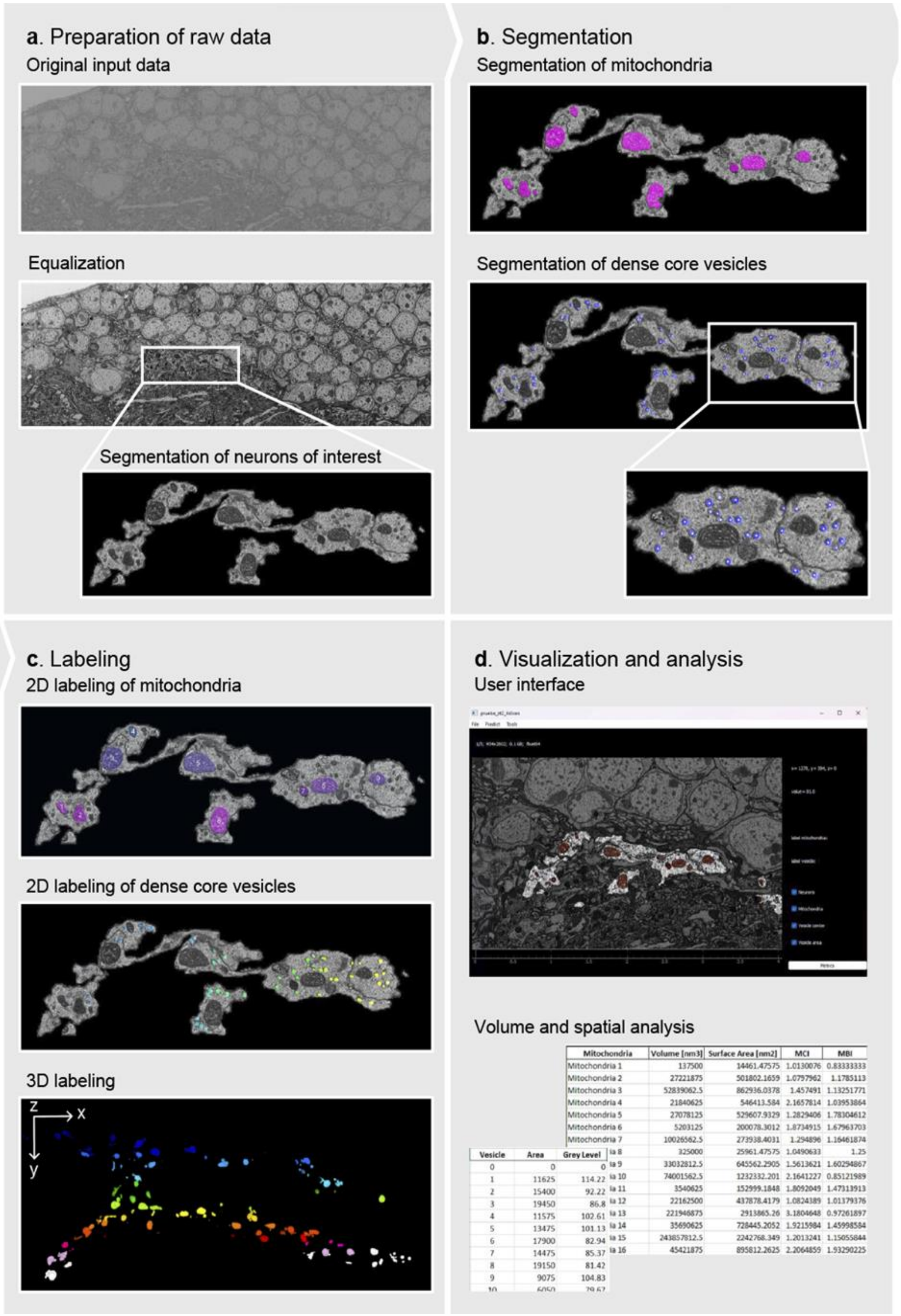


**Supplementary Figure 2. Overview of the algorithm developed for neuron and organelle segmentation. a**, Preparation of raw data, including equalization and segmentation of neurons of interest. **b**, Segmentation of mitochondria and dense core vesicles using deep learning models. **c**, 2D and 3D labeling of segmented mitochondria and vesicles. **d**, Visualization of the results of the segmentation and subsequent volume and spatial analysis through the user interface.

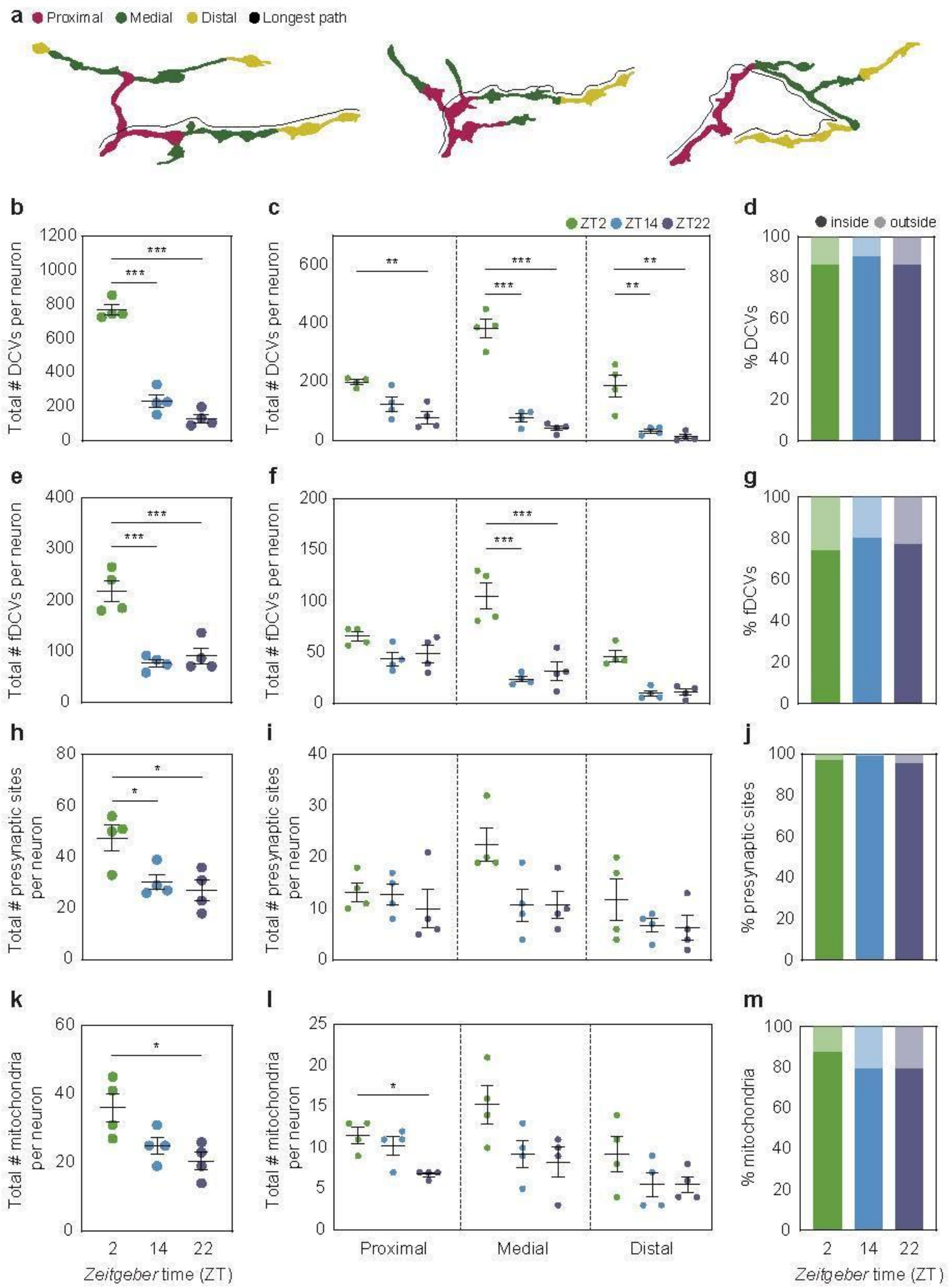


**Supplementary Figure 3. Distribution of free and fused DCVs, presynaptic sites and mitochondria. a**, Schematic diagram of representative neurites taken from the ZT2 (left), 14 (middle) and 22 (right) volumes showing the proximal (magenta), medial (green) and distal (yellow) regions. The longest neurite in each neuronal terminal was divided in thirds and those lengths were applied to the remaining neurites. The distribution in proximal, medial or distal regions refers to objects segmented in the initial, subsequent and terminal thirds of each neurite. **b, e, h, k** Total (including the objects located outside of varicosities) number of free DCVs (**b**), fDCVs (**e**), PSs (**h**) and mitochondria (**k**) at the different time points. **c**, **f**, **i**, **l** Quantity and distribution within the proximal, medial and distal portions of the neurites of free DCVs (**c**), fDCVs (**f**), PSs (**i**) and mitochondria (**l**). **d**, **g**, **j**, **k** Proportion of each of the objects exhibited within or outside of varicosities at each time point. The panel displayed in **k** corresponds to the data already shown in **Figure 3b**, and it has been included herein to enable direct comparison. In all graphs, error bars indicate the standard error of the mean (SEM). Asterisks indicate statistically significant differences: * p < 0.05, ** p < 0.01, *** p < 0.001. Non-significant differences are not shown. Details can be found in **Supplementary Table 3**.

| Experiment | Group | n | Analysis of deviance | | | Contrasts | | |
|---|---|---|---|---|---|---|---|---|
| | | | Chisq. | df | p-value | vs. | z | p-value |
| # DCVs proximal (Figure 2c) | ZT2 | 21 | 13,504 | 2 | 0,001 | 2 vs 14 | 2,16 | 0.092 |
| | ZT14 | 22 | | | | 2 vs 22 | 3,576 | 0.001 |
| | ZT22 | 18 | | | | 14 vs 22 | 1,582 | 0.341 |
| # DCVs medial (Figure 2c) | ZT2 | 28 | 117,84 | 2 | < 0.001 | 2 vs 14 | 6,382 | < 0.001 |
| | ZT14 | 18 | | | | 2 vs 22 | 9,763 | < 0.001 |
| | ZT22 | 19 | | | | 14 vs 22 | 3,926 | < 0.001 |
| # DCVs distal (Figure 2c) | ZT2 | 14 | 90,512 | 2 | < 0.001 | 2 vs 14 | 4,972 | < 0.001 |
| | ZT14 | 10 | | | | 2 vs 22 | 8,878 | < 0.001 |
| | ZT22 | 14 | | | | 14 vs 22 | 3,763 | < 0.001 |
| # fDCVs proximal (Figure 2f) | ZT2 | 21 | 3,2742 | 2 | 0,195 | 2 vs 14 | 1,76 | 0.235 |
| | ZT14 | 21 | | | | 2 vs 22 | 0,385 | 1 |
| | ZT22 | 18 | | | | 14 vs 22 | -1,313 | 0.568 |
| # fDCVs medial (Figure 2f) | ZT2 | 29 | 50,08 | 2 | > 0.001 | 2 vs 14 | 5,997 | < 0.001 |
| | ZT14 | 17 | | | | 2 vs 22 | 4,895 | < 0.001 |
| | ZT22 | 19 | | | | 14 vs 22 | -1,557 | 0.358 |
| # fDCVs distal (Figure 2f) | ZT2 | 14 | 33,463 | 2 | > 0.001 | 2 vs 14 | 3,968 | < 0.001 |
| | ZT14 | 10 | | | | 2 vs 22 | 4,989 | < 0.001 |
| | ZT22 | 14 | | | | 14 vs 22 | 0,644 | 1 |
| # PSs proximal (Figure 3c) | ZT2 | 21 | 0,6143 | 2 | 0,736 | 2 vs 14 | 0,235 | 1 |
| | ZT14 | 22 | | | | 2 vs 22 | 0,772 | 1 |
| | ZT22 | 18 | | | | 14 vs 22 | 0,557 | 1 |

| | | | | | | | | |
|---|---|---|---|---|---|---|---|---|
| # PSs medial (Figure 3c) | ZT2 | 29 | 2,9345 | 2 | 0,231 | 2 vs 14 | 1,608 | 0.323 |
| | ZT14 | 18 | | | | 2 vs 22 | 2,004 | 0.135 |
| | ZT22 | 20 | | | | 14 vs 22 | 0,351 | 1 |
| # PSs distal (Figure 3c) | ZT2 | 14 | 5,5653 | 2 | 0,062 | 2 vs 14 | 0,727 | 1 |
| | ZT14 | 10 | | | | 2 vs 22 | 2,356 | 0.056 |
| | ZT22 | 14 | | | | 14 vs 22 | 1,474 | 0.421 |

**Supplementary Table 1. Statistical parameters for generalized linear models**. **n**: sample size **Chisq**: chi squared. **df**: degrees of freedom. **vs**: compared groups.

| Experiment | Group | n | ANOVA | | Contrasts | | | |
|---|---|---|---|---|---|---|---|---|
| | | | F-value | p-value | vs. | t | df | p-value |
| GRIP (Figure 2j) | ZT2 | 28 | F(0.05;2;66)= 8.776 | > 0.001 | 2 vs 14 | 3.932 | 66 | < 0.001 |
| | ZT14 | 22 | | | 2 vs 22 | 2.126 | 66 | 0.112 |
| | ZT22 | 20 | | | 14 vs 22 | -2.411 | 66 | 0.056 |
| MCI vs ZT (Figure 4c) | ZT2 | 144 | F(0.05;2;321,03) = 11.726 | > 0.001 | 2 vs 14 | -3.511 | 321 | 0,002 |
| | ZT14 | 100 | | | 2 vs 22 | -4.425 | 321 | <0.001 |
| | ZT22 | 82 | | | 14 vs 22 | -1.040 | 320 | 0,898 |
| Distance between varicosities (Figure 5c) | ZT2 | 62 | F(0.05;2,161)= 0.506 | 0,604 | 2 vs 14 | -0.037 | 161 | 1 |
| | ZT14 | 49 | | | 2 vs 22 | -0.912 | 159 | 1 |
| | ZT22 | 53 | | | 14 vs 22 | -0.823 | 160 | 1 |
| Girth (Figure 5d) | ZT2 | 64 | F(0.05;2,163)= 10.672 | > 0.001 | 2 vs 14 | -1.304 | 163 | 0.582 |
| | ZT14 | 50 | | | 2 vs 22 | 3.380 | 161 | 0.003 |
| | ZT22 | 52 | | | 14 vs 22 | 4.436 | 161 | < 0.001 |

**Supplementary Table 2. Statistical parameters for general linear models**. **n**: sample size. **vs**: compared groups. **df**: degrees of freedom.

| Experiment | Group | ANOVA | | Contrast | | | |
|---|---|---|---|---|---|---|---|
| | | F-value | p-value | vs. | t | df | p-value |

| | | | | | | | |
|---|---|---|---|---|---|---|---|
| Neuron volume (Figure 1f) | ZT2 | F(0.05;2;9)= 12.151 | 0,003 | 2 vs 14 | -0,677 | 9 | 1 |
| | ZT14 | | | 2 vs 22 | 4,567 | 9 | 0.004 |
| | ZT22 | | | 14 vs 22 | 3,891 | 9 | 0.011 |
| DCV density per neuron (Figure 2b) | ZT2 | F(0.05;2;9)= 62.211 | < 0.001 | 2 vs 14 | -9,386 | 9 | < 0.001 |
| | ZT14 | | | 2 vs 22 | 9,912 | 9 | < 0.001 |
| | ZT22 | | | 14 vs 22 | 0,526 | 9 | 1 |
| fDCV density per neuron (Figure 2e) | ZT2 | F(0.05;2;9)= 30.645 | < 0.001 | 2 vs 14 | -7,828 | 9 | < 0.001 |
| | ZT14 | | | 2 vs 22 | 3,806 | 9 | 0.013 |
| | ZT22 | | | 14 vs 22 | -4,022 | 9 | 0.009 |
| PSs density per neuron (Figure 3d) | ZT2 | F(0.05;2;9)= 3.948 | 0,059 | 2 vs 14 | -2,589 | 9 | 0.088 |
| | ZT14 | | | 2 vs 22 | 0,348 | 9 | 1 |
| | ZT22 | | | 14 vs 22 | -2,241 | 9 | 0.155 |
| Mitochondrial volume (Figure 4b) | ZT2 | F(0.05;2;9)= 2.342 | 0,152 | 2 vs 14 | -1,148 | 9 | 0.840 |
| | ZT14 | | | 2 vs 22 | 2,163 | 9 | 0.180 |
| | ZT22 | | | 14 vs 22 | 1,015 | 9 | 1 |
| Total # DCVs (Suppl. Figure 3b) | ZT2 | F(0.05;2;9)= 127.523 | < 0.001 | 2 vs 14 | -12,484 | 9 | < 0.001 |
| | ZT14 | | | 2 vs 22 | 14,867 | 9 | < 0.001 |
| | ZT22 | | | 14 vs 22 | 2,383 | 9 | 0.120 |
| # DCVs per neuron proximal (Suppl. Figure 3c) | ZT2 | F(0.05;2;9)= 10.410 | 0,005 | 2 vs 14 | -2,800 | 9 | 0,062 |
| | ZT14 | | | 2 vs 22 | 4,520 | 9 | 0,004 |
| | ZT22 | | | 14 vs 22 | 1,721 | 9 | 0,358 |
| # DCVs per neuron medial (Suppl. Figure 3c) | ZT2 | F(0.05;2;9)= 89.278 | < 0.001 | 2 vs 14 | -10,888 | 9 | < 0.001 |
| | ZT14 | | | 2 vs 22 | 12,152 | 9 | < 0.001 |
| | ZT22 | | | 14 vs 22 | 1,264 | 9 | 0,713 |
| # DCVs per neuron distal (Suppl. Figure 3c) | ZT2 | F(0.05;2;9)= 17.182 | < 0.001 | 2 vs 14 | -4,735 | 9 | 0,003 |
| | ZT14 | | | 2 vs 22 | 5,361 | 9 | 0,001 |
| | ZT22 | | | 14 vs 22 | 0,626 | 9 | 1 |
| Total # fDCVs | ZT2 | | < 0.001 | 2 vs 14 | -6,347 | 9 | < 0.001 |

| | | | | | | | |
|---|---|---|---|---|---|---|---|
| (Suppl. Figure 3e) | ZT14 | F(0.05;2;9)= 24.441 | | 2 vs 22 | 5,713 | 9 | < 0.001 |
| | ZT22 | | | 14 vs 22 | -0,635 | 9 | 1 |
| # fDCV per neuron proximal (Suppl. Figure 3f) | ZT2 | F(0.05;2;9)= 3.226 | 0,088 | 2 vs 14 | -2,421 | 9 | 0,116 |
| | ZT14 | | | 2 vs 22 | 1,877 | 9 | 0,28 |
| | ZT22 | | | 14 vs 22 | -0,544 | 9 | 1 |
| # fDCV per neuron medial (Suppl. Figure 3f) | ZT2 | F(0.05;2;9)= 23.940 | < 0.001 | 2 vs 14 | -6,269 | 9 | < 0.001 |
| | ZT14 | | | 2 vs 22 | 5,671 | 9 | < 0.001 |
| | ZT22 | | | 14 vs 22 | -0,598 | 9 | 1 |
| # fDCV per neuron distal (Suppl. Figure 3f) | ZT2 | F(0.05;2;9)= 28.533 | < 0.001 | 2 vs 14 | -6,653 | 9 | < 0.001 |
| | ZT14 | | | 2 vs 22 | 6,425 | 9 | < 0.001 |
| | ZT22 | | | 14 vs 22 | -0,228 | 9 | 1 |
| Total # PSs (Suppl. Figure 3h) | ZT2 | F(0.05;2;9)= 7.263 | 0,013 | 2 vs 14 | -2,983 | 9 | 0.046 |
| | ZT14 | | | 2 vs 22 | 3,546 | 9 | 0.019 |
| | ZT22 | | | 14 vs 22 | 0,562 | 9 | 1 |
| # PSs per neuron proximal (Suppl. Figure 3i) | ZT2 | F(0.05;2;9)= 0.435 | 0,66 | 2 vs 14 | -0,133 | 9 | 1 |
| | ZT14 | | | 2 vs 22 | 0,866 | 9 | 1 |
| | ZT22 | | | 14 vs 22 | 0,733 | 9 | 1 |
| # PSs per neuron medial (Suppl. Figure 3i) | ZT2 | F(0.05;2;9)= 5.235 | 0,031 | 2 vs 14 | -2,802 | 9 | 0,062 |
| | ZT14 | | | 2 vs 22 | 2,802 | 9 | 0,062 |
| | ZT22 | | | 14 vs 22 | 0,000 | 9 | 1 |
| # PSs per neuron distal (Suppl. Figure 3i) | ZT2 | F(0.05;2;9)= 1.197 | 0,346 | 2 vs 14 | -1,272 | 9 | 0,706 |
| | ZT14 | | | 2 vs 22 | 1,399 | 9 | 0,586 |
| | ZT22 | | | 14 vs 22 | 0,127 | 9 | 1 |
| Total # mitochondria (Suppl. Figure 3k) | ZT2 | F(0.05;2;9)= 6.271 | 0,020 | 2 vs 14 | -2,443 | 9 | 0.112 |
| | ZT14 | | | 2 vs 22 | 3,442 | 9 | 0.022 |
| | ZT22 | | | 14 vs 22 | 0,999 | 9 | 1 |
| | ZT2 | | 0,009 | 2 vs 14 | -1,030 | 9 | 0,989 |

| | | | | | | | |
|---|---|---|---|---|---|---|---|
| # mitochondria per neuron proximal (Suppl. Figure 3l) | ZT14 | F(0.05;2;9)= 8.236 | | 2 vs 22 | 3,915 | 9 | 0,011 |
| | ZT22 | | | 14 vs 22 | 2,885 | 9 | 0,054 |
| # mitochondria per neuron medial (Suppl. Figure 3l) | ZT2 | F(0.05;2;9)= 3.844 | 0,062 | 2 vs 14 | -2,197 | 9 | 0,167 |
| | ZT14 | | | 2 vs 22 | 2,563 | 9 | 0,092 |
| | ZT22 | | | 14 vs 22 | 0,366 | 9 | 1 |
| # mitochondria per neuron distal (Suppl. Figure 3l) | ZT2 | F(0.05;2;9)= 1.819 | 0,217 | 2 vs 14 | -1,652 | 9 | 0,399 |
| | ZT14 | | | 2 vs 22 | 1,652 | 9 | 0,399 |
| | ZT22 | | | 14 vs 22 | 0,000 | 9 | 1 |
| Total # varicosities (Figure 5b) | ZT2 | F(0.05;2;9)= 3.146 | 0,092 | 2 vs 14 | -2,319 | 9 | 0.140 |
| | ZT14 | | | 2 vs 22 | 1,988 | 9 | 0.230 |
| | ZT22 | | | 14 vs 22 | -0,331 | 9 | 1 |
| # varicosities per neuron proximal (Figure 5b) | ZT2 | F(0.05;2;9)= 0.470 | 0,64 | 2 vs 14 | 0,233 | 9 | 1 |
| | ZT14 | | | 2 vs 22 | 0,699 | 9 | 1 |
| | ZT22 | | | 14 vs 22 | 0,931 | 9 | 1 |
| # varicosities per neuron medial (Figure 5b) | ZT2 | F(0.05;2;9)= 2.433 | 0,143 | 2 vs 14 | -2,071 | 9 | 0.200 |
| | ZT14 | | | 2 vs 22 | 1,694 | 9 | 0.373 |
| | ZT22 | | | 14 vs 22 | -0,376 | 9 | 1 |
| # varicosities per neuron distal (Figure 5b) | ZT2 | F(0.05;2;9)= 0.632 | 0,554 | 2 vs 14 | -0,973 | 9 | 1 |
| | ZT14 | | | 2 vs 22 | 0,000 | 9 | 1 |
| | ZT22 | | | 14 vs 22 | -0,973 | 9 | 1 |

**Supplementary Table 3. Statistical parameters for general linear models with n = 4**. **vs**: compared groups. **df**: degrees of freedom.
